# Stereospecific rapid activation of Transcription Factor EB (TFEB) by Levacetylleucine (NALL)

**DOI:** 10.1101/2025.06.06.657607

**Authors:** Lianne C. Davis, Rebecca Brain, Grant Churchill, Mallory Factor, Taylor Fields, Frances Platt, Marc Patterson, Michael Strupp, Antony Galione

## Abstract

N-acetyl L-leucine (NALL, USAN or levacetylleucine, INN or trade name Aqneursa™) is an FDA-approved agent for the treatment of Niemann-Pick disease type C (NPC). The N-acetyl group renders the compound a prodrug of L-leucine, making it a substrate for membrane-spanning monocarboxylate transporters (MCTs), which are ubiquitously expressed delivering NALL to all tissues with high capacity, including to the central nervous system. NALL enters enzyme-controlled pathways that correct metabolic dysfunction and enhance mitochondrial bioenergetics. Because NALL improves energy production (adenosine triphosphate, ATP) and ameliorates lysosomal function, it is potentially a therapy for a broad range of neurodegenerative and neurodevelopmental disorders (in addition to lysosomal storage disorders) in which energy homeostasis and lysosomal function are impaired.

Here, we have performed a series of *in vitro* studies which reveal an additional aspect of NALL’s polypharmacological mechanism of action. The studies demonstrate a direct lysosomal effect whereby NALL rapidly activates the translocation of the Transcription Factor EB (TFEB, a master regulator of lysosomal biogenesis and autophagy) from the cytoplasm to the nucleus in HeLa cells. The activation of TFEB is known to trigger the activation of specific genes that restore lysosomal biogenesis and function, as well as autophagy. Consistent with this, we show that NALL increases production of a TFEB target gene LAMP1, an integral lysosomal membrane protein responsible for maintaining lysosomal integrity, function and pH. This *in vitro* effect occurs at concentrations consistent with concentrations in plasma in humans after standard therapeutic dosing. We further demonstrated that acetylation is critical to this aspect of NALL’s mechanism of action, as L-leucine itself had no effect on the activation of TFEB. Consistent with previous studies N-acetyl-D-leucine was inactive and also had no effect. Similarly, N-acetyl-DL-leucine also had only a modest effect, providing further evidence that N-acetyl-D-leucine is even antagonistic and inhibits the effects of the active L-enantiomer.

This mechanism of action of NALL to activate TFEB signalling, thereby enhancing lysosomal and autophagic function, further elucidates the ways by which this compound targets the fundamental etiology of rare and common neurodegenerative disorders, from Niemann-Pick disease type C to Parkinson’s disease. Based on its mechanism of action by improving the mitochondrial-lysosomal axis, NALL has the potential to be an effective therapy for a broad range of neurological and neurodevelopment conditions.

## Introduction

Levacetylleucine (N-acetyl-L-leucine, NALL) was approved by the US Food and Drug Administration (FDA) as a monotherapy for the treatment of the rare genetic lysosomal storage disease (LSD) Niemann-Pick disease type C (NPC) (Beninger, 2024; Mullard, 2024; van Gool et al., 2025) on 24^th^ September 2024. Though NPC itself is an ultra-rare disease affecting approximately 1/100,000 live births (Tifft, 2024), it is 1 of over 70 rare monogenic diseases classified as lysosomal storage disorders (LSD) that collectively affect 1:5000 live births (Platt et al., 2018). NPC is caused by mutations in *NPC1* (95% of cases) or *NPC2* (5% of cases) genes that encode proteins important for intracellular trafficking and transport of cholesterol and sphingolipids from the late endosome/lysosome (Alavi et al., 2013; Fusco et al., 2013; Naureckiene et al., 2000; Park et al., 2003; Steinberg et al., 1994). NPC1 or NPC2 dysfunction leads to the accumulation LDL-derived cholesterol and sphingolipids in the late endocytic system leading to lysosomal enlargement, disruption of intracellular lipid trafficking and autophagy, dysregulation of the mTOR signalling pathway, reduction in lysosomal Ca^2+^ levels and release, perturbation of cellular metabolism and ultimately cell death (Platt et al., 2018), and dysregulation of membrane contact sites (MCS) between the ER and lysosomes and gain of mitochondrial-lysosomal MCS (Hoglinger et al., 2019).

Although NPC affects all organs, the central nervous system (CNS) is particularly vulnerable, and the combined effects of the biochemical and cell biological defects in NPC lead to widespread pathology in the brain, driving and exacerbating neuroinflammation and neurodegeneration (Walterfang et al., 2020). The clinical manifestations of this are seen in the broad range of neurological signs and symptoms patients with NPC (colloquially referred to as “childhood Alzheimer’s”) suffer from, including gait abnormalities, tremors, ataxia, dystonia, speech impairment, seizures, dysphagia, motor dysfunction, progressive cognitive impairment, and dementia (Patterson et al., 2020).

NALL (Aqneursa^TM^) is an orally administered amino-acid derivative. The N-acetylation makes it a substrate for membrane-spanning monocarboxylate transporters (MCTs), bypassing LAT (the leucine transporter) thereby delivering higher levels of NALL to the cytosol. MCTs are a family with 14 members and are ubiquitously expressed throughout the body, enabling NALL’s delivery to all tissues, including the central nervous system by readily crossing the blood-brain barrier (Churchill et al., 2021). NALL enters enzyme-controlled pathways that correct metabolic dysfunction and enhance energy (ATP) production which also leads to an improvement in lysosomal function (Kaya et al., 2020; Kaya et al., 2021). This has multiple knock-on effects: mitochondrial and lysosomal function are intrinsically linked, and have been shown to even physically interact at membrane contact sites (Martello et al., 2020). NALL’s normalization of mitochondrial function and energy metabolism improves lysosomal function, leading to a reduction in the storage of unesterified cholesterol and sphingolipids (Kaya et al., 2021).

NALL has been demonstrated in various animal models to restore neuronal function and substantially delay and slow neurodegeneration as well as dampen neuroinflammation, consistent with the drug’s neuroprotective effects (Hegdekar et al., 2021; Kaya et al., 2021). Consistent with these pre-clinical data, multiple clinical studies have robustly demonstrated that NALL rapidly improves neurological signs, symptoms, and function, but also has a long-term disease-modifying, neuroprotective effect in NPC (Bremova-Ertl et al., 2022; Bremova-Ertl et al., 2024). NALL has also demonstrated the potential to be effective for a broad range of neurodegenerative and neurodevelopmental disorders, including other lysosomal diseases such as GM2 gangliosidosis and Batten’s Disease, inherited cerebellar ataxias, and common diseases including Parkinson’s Disease and Dementias (Cortes and La Spada, 2019).

To further explore the pathways by which NALL targets fundamental disease mechanisms, we conducted a series of *in vitro* experiments to examine its role in modulating Transcription Factor EB (TFEB) activity. TFEB, a key transcription factor that impacts lysosomes, orchestrates the expression of genes (the coordinated lysosomal expression and regulation (CLEAR) network) involved in lysosomal biogenesis and autophagy. It promotes the expression of genes required for autophagosome formation, lysosome biogenesis, and lysosomal function and exocytosis (Napolitano and Ballabio, 2016). TFEB contains basic helix-loop-helix-leucine zipper domains (bHLH-Zip) and belongs to the microphthalmia MIT-TFE family of transcription factors (Puertollano et al., 2018), and is highly expressed in CNS (Cortes and La Spada, 2019). In its inactive phosphorylated state, TFEB resides in the cytoplasm; upon dephosphorylation it dissociates from 14-3-3 proteins and translocates to the nucleus, to regulate gene expression (Rosenbaum and Maxfield, 2011). Dysregulation of TFEB activity is implicated in various neurodegenerative and neurodevelopmental diseases, including NPC (Chen et al., 2024). Previous studies have demonstrated that targeting the TFEB pathway has neuroprotective effects in various *in vivo* or *in vitro* models of Alzheimer’s disease (Gu et al., 2022), and it has recently been demonstrated that activation of TFEB is beneficial and promotes lysosomal clearance in *NPC1*^-/-^ cells

In these studies, we show in wild type Hela cells that NALL stimulates TFEB translocation to the nucleus, which would serve to enhance lysosomal biogenesis, function, and autophagic flux, and may therefore be potentially beneficial in neurodegenerative diseases (Gu et al., 2022).This effect provides further evidence of NALL’s novel mechanism of action in addition to improving mitochondrial bioenergetics (Kaya et al., 2021).

## Methods

### Chemicals

N-acetyl L-leucine (NALL) was obtained from IntraBio, Inc, 201 W 5th Street, Suite 1100, Austin, TX 78701, USA. NALL was solubilised directly into DMEM at 2 mM, and adjusted to pH 7.4 using NaOH.

### Cell Culture and Transfection

HeLa cells were cultured in DMEM supplemented with 10% v/v FCS, 2 mM glutamine, 100 U/ml penicillin and 100 µg/ml streptomycin, at 37^°^C under 5% CO_2_. Cells were trypsinised and seeded onto CellView Slides (Greiner Bio-One).

One to two days after sub-culturing, cells were transiently transfected with JetPEI reagent in a 5:2 ratio with DNA. Per well, cells were transfected with 100 ng TFEB tagged on its C-terminus with either EGFP (Addgene plasmid # 38119) or mScarlet3 (produced in-house) for 4-6 h. Transfection medium was removed and replaced with fresh DMEM with or without agents and incubated overnight at 37°C. The next day, TFEB-EGFP-expressing cells were loaded for 1 h at 37^°^C with NucSpot® Live 650 Nuclear Stain (Biotium) in the continued presence of drugs as required. Cells were then transferred into extracellular medium (ECM, mM: 121 NaCl, 5.4 KCl, 0.8 MgCl_2_, 1.8 CaCl_2_, 6 NaHCO_3_, 25 HEPES, 10 glucose) that maintained the overnight treatment reagents and these live cells were imaged immediately.

Cells expressing TFEB-mScarlet3 were treated with similar reagent protocols but could be fixed with 4% PFA (in PBS) after reagent treatments. After fixation and permeabilization (0.1% Triton X-100 in PBS for 15 min), nuclei were labelled with NucSpot® Live 488 Nuclear Stain (Biotium) for 10 min in PBS and imaged within 2 days.

### Microscopy

Cells were imaged at room temperature using a Nikon A1R laser-scanning confocal equipped with a Plan ApoVC 20x DIC N2 (NA: 0.75) or Plan Fluor 40x oil DIC H N2 (NA: 1.3) objective. In Channel-Series mode, green, red or far-red fluorophores were alternately excited (ex/em): 488/525 nm, 561/595 nm, and 640/700 nm respectively.

### Nuclear Translocation Analysis

To quantify the translocation of fluorescent TFEB from the cytoplasm to the nucleus, we analysed the Pearson’s Correlation Coefficient between TFEB and the orthogonal nuclear stain in single cells. Cells with a cytoplasmic location have negative coefficients, whereas nuclear translocation is reflected by positive coefficients. Cells with nuclear TFEB (partial or completely translocated) were defined as having a Pearson’s coefficient >0, and the number of cells satisfying this criterion expressed as a percentage of the total number.

### In-Cell Western for detection of LAMP1

HeLa were seeded into a 96-well flat-bottom μclear black plate (Greiner Bio-one). Following treatment with agents, cells were fixed with 4% PFA and permeabilised with 0.1% Triton X-100 for 10 min. Following block in Odyssey blocking buffer (LICORbio), cells were incubated with mouse anti-LAMP1 antibody (H4A3, DSHB at The University of Iowa) and was detected using IRDye 800CW goat anti-mouse IgG secondary antibody (LICORbio). Cells were co-stained with CellTag 700 Stain for In-Cell Western Assays (LICORbio) for accurate normalization to cell number. Cells were scanned using an Odyssey M Imaging System in the 700 and 800 nm channels and analysed using Empiria Studio (LICORbio).

### Reverse Transcription Quantitative PCR (RT-qPCR)

Total RNA was extracted from HeLa cells using a RNeasy Plus Mini kit (Qiagen) following the manufacturer’s instructions. cDNA was prepared using an iScript cDNA synthesis kit in a MyCycler Thermal Cycler system (both Bio-Rad). qPCR was then performed with a CFX96 Real-Time PCR instrument (Bio-Rad) using PowerUp SYBR Green (Applied Biosystems) and the primer pairs specified in **Table 1**. Threshold cycle (C_t_) values were normalised to the reference gene ß-actin using the comparative C_t_ method, on a scale where beta-actin expression equals 10,000 units.

**Table 1.**
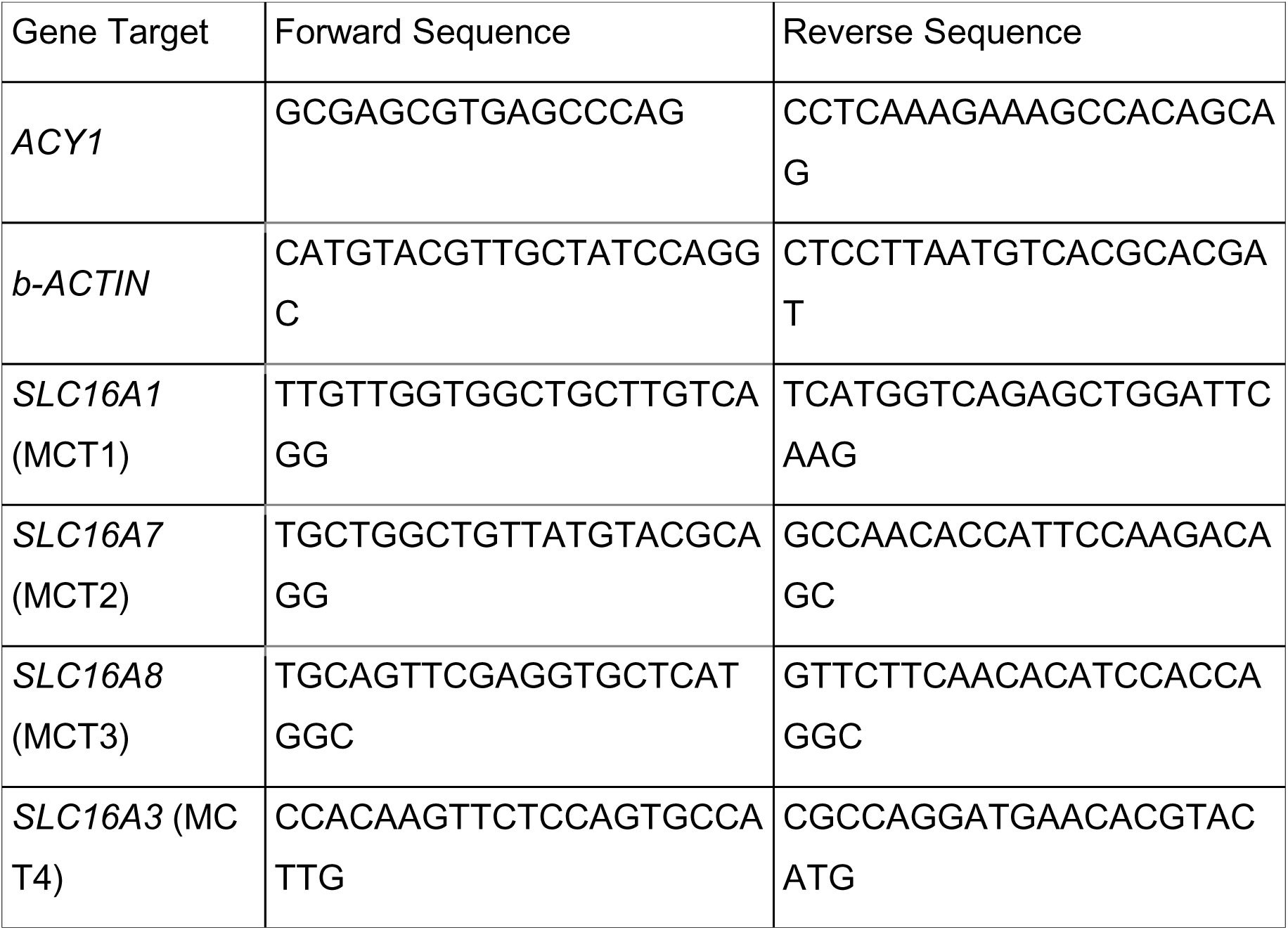
Primer sequences for RT-qPCR, sourced from OriGene and PrimerBank.

## Results

We have previously demonstrated that NALL acts as a pro-drug and enters cells via MCT transporters to generate high concentrations of cytosolic L-leucine in cells (Churchill et al., 2021). We first investigated whether HeLa cells would be a suitable model system which expresses the key components of the NALL transport and metabolism mechanisms and found that HeLa cells not only express the monocarboxylate transporters (MCT1, MCT2, MCT3, MCT4) but also the aminoacylase 1 enzyme (ACY1), as determined by RT-qPCR (primer sequences in **Table 1**). Expression was normalised to β-actin, on a scale where β-actin expression equals 10,000 arbitrary units. We found that HeLa cells contain mRNA for several monocarboxylate transporter types, including MCT1, (**Fig. S1**) as well as an acylase shown to hydrolyze N-acetyl groups from amino acids (Bloch and Borek, 1946; Bloch and Rittenberg, 1947). The resulting L-leucine enters metabolic pathways to increase ATP synthesis, but also acts to clear stored lipids from lysosomes (Kaya et al., 2020).

To test for an effect of the drug on lysosomal function, we examined the effect of treating HeLa with NALL on TFEB, a master transcription factor that regulates lysosome biogenesis. A hallmark of its activation is the translocation of TFEB from the cytoplasm to the nucleus (Settembre et al., 2012). We found that incubation of cells overnight with extracellular NALL caused a concentration-dependent appearance of TFEB-GFP in the nucleus, with activity in the high micromolar range (**Fig. 1A-C**). Curve fitting of the data plotted as either Pearson’s coefficient or percentage of cells with nuclear TFEB, calculated an EC_50_ of 225 μM and 276 μM, respectively. This concentration range coincides with plasma levels observed in mice dosed with therapeutic concentrations of NALL with a peak serum concentration (C_max_) of 100-200 µM (Churchill et al., 2020). Indeed, at 100 μM, the Pearson’s coefficient was 28.4% of the maximum response and the number of cells with nuclear TFEB was 59.5% of the maximum.

**Figure 1.**
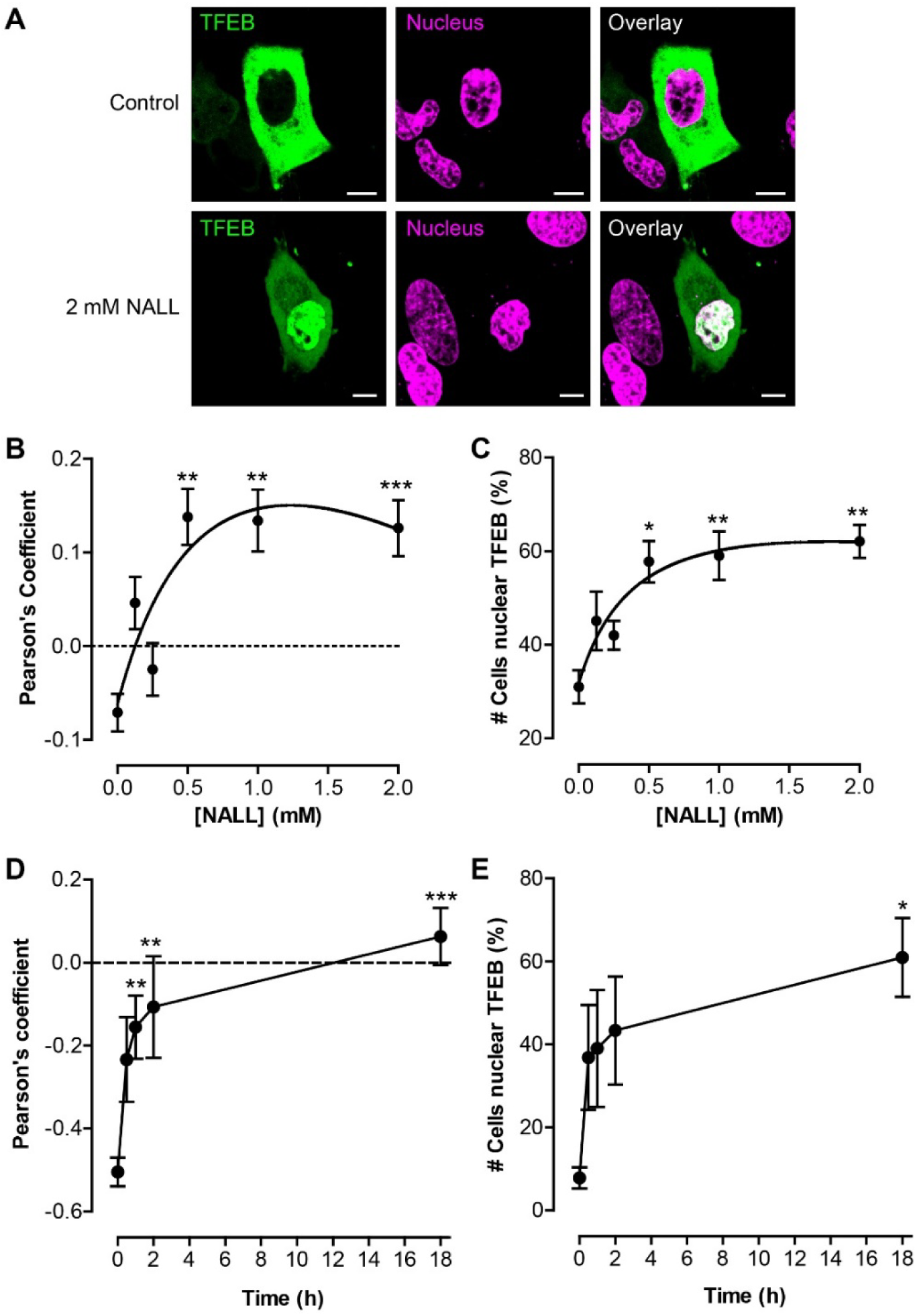
Levacetylleucine induces nuclear translocation of TFEB. HeLa cells were transfected with TFEB-EGFP (A-C) or TFEB-mScarlet3 (D,E) and then incubated with levacetylleucine (NALL) at different concentrations or times. Cells were then counter-stained with the NucSpot 488 (D,E) or NucSpot 650 (A-C) to label the nucleus and TFEB translocation quantified by colocalization with NucSpot. (A) Single-cell images of TFEB-EGFP (green) and NucSpot 650 (magenta). (B,C) Concentration-response of HeLa treated for 18 hours with NALL. Nuclear localization is plotted as either the Pearson’s coefficient (B) or derived as the percentage of cells with nuclear TFEB (C). (D,E) Time course of the effect of 2 mM NALL upon TFEB, either expressed as the Pearson’s coefficient (D) or the percentage of cells with nuclear TFEB (E). The dotted lines (B,D) highlight a correlation coefficient of zero used for thresholding the percentage of nuclear cells. Data are expressed as the mean ± SEM of 179-215 cells, and significance determined by ANOVA test, with significance depicted as * *p* < 0.05, ** *p* < 0.01, or *** *p* < 0.001 compared to respective control, 0 mM (B,C) or t = 0 (D,E).

The degree of TFEB translocation at different times after NALL addition was quantified. There was a rapid translocation, with a half-maximal effect seen within 60 min (**Fig. 1D, E**), showing that NALL acts rapidly.

Importantly, the pharmacological concentration-response provided by NALL was not shared by non-acetylated L-leucine over a similar concentration range (**Fig. 2**), which is the major metabolite of NALL through the action of amino acid acylases (Birnbaum et al., 1952), demonstrating that this effect required the acetylation of leucine as in NALL. This effect may be transient since NALL is converted to L-leucine by cellular acylases (Bloch and Rittenberg, 1947; Churchill et al., 2020).

**Figure 2.**
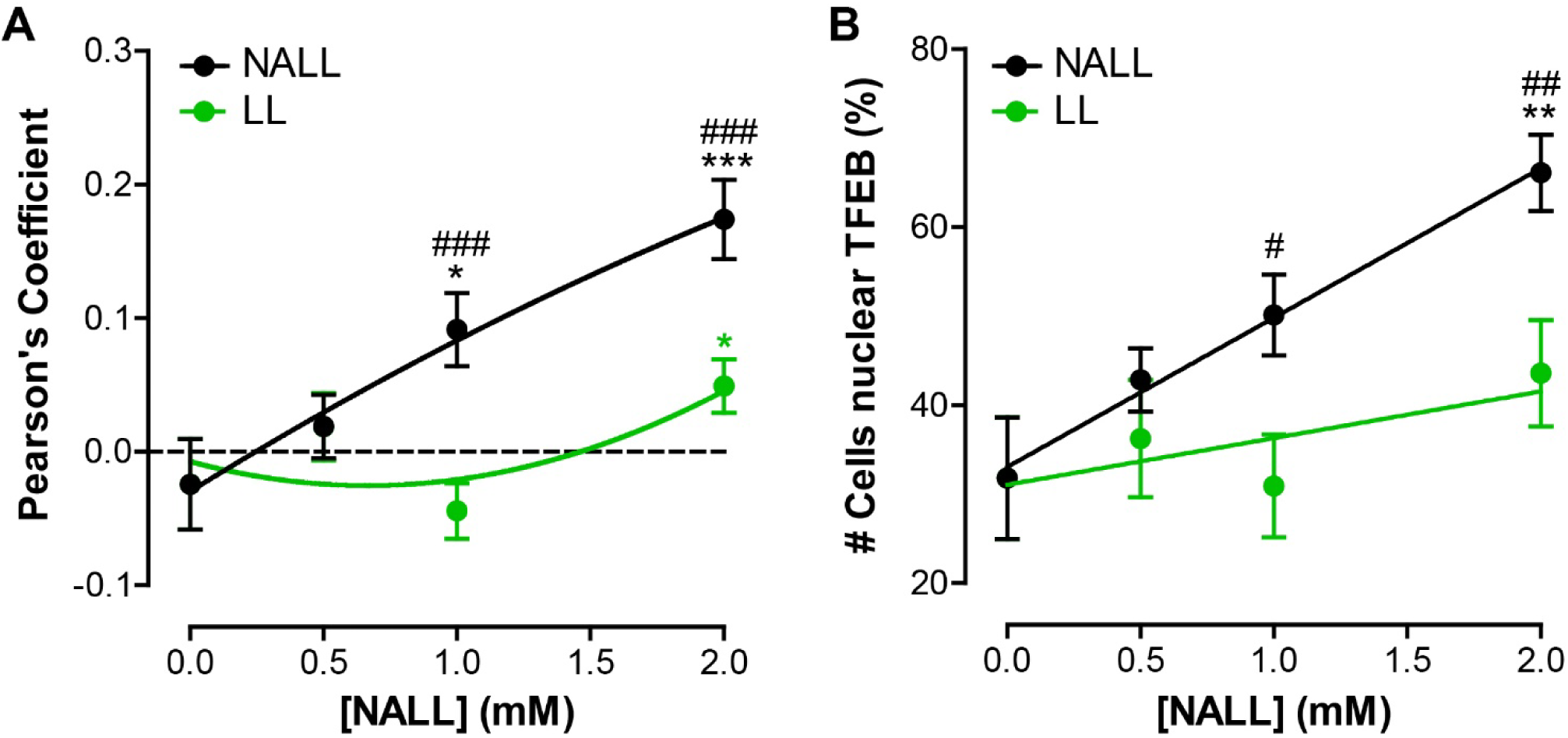
Levacetylleucine induces TFEB translocation more than does L-leucine. TFEB-EGFP translocation in HeLa cells incubated for 18 hours with different concentrations of either levacetylleucine (NALL, black) or L-Leucine (LL, green). Translocation is expressed either as the Pearson’s correlation coefficient with NucSpot 650 (A) or as the percentage of cells with nuclear TFEB (B). Data are expressed as the mean ± SEM of 104-161 cells, and significance determined by nonparametric ANOVA test (A) or unpaired t-test (B), with significance depicted as * *p* < 0.05, ** *p* < 0.01, or *** *p* < 0.001 compared to the vehicle control, i.e. 0 mM. Comparing NALL and LL: #, P<0.05; ###, P < 0.001.

Previously, a racemic mixture of the D- and L-forms-, (N-acetyl D-L-leucine, Tanganil) has also been observed to have beneficial effects in lysosomal storage diseases (Kaya et al., 2020). Therefore, we compared different enantiomers for their ability to drive TFEB translocation, either the separate enantiomers NALL and N-acetyl D-leucine or the racemate N-acetyl D-L-leucine (**Fig. 3**). Whilst NALL caused a robust translocation of TFEB into the nucleus, the D-enantiomer, N-acetyl D-leucine, was without effect over this time period. This confirms the stereochemistry is crucial for the effect of NALL. Moreover, the racemate mixture of both (N-acetyl D-L-leucine) also failed to drive TFEB translocation, indicating that the presence of the D-enantiomer in the racemate antagonizes the effect of the L-enantiomer, NALL. In this assay then, the NALL is the superior enantiomer for inducing TFEB activation.

**Figure 3.**
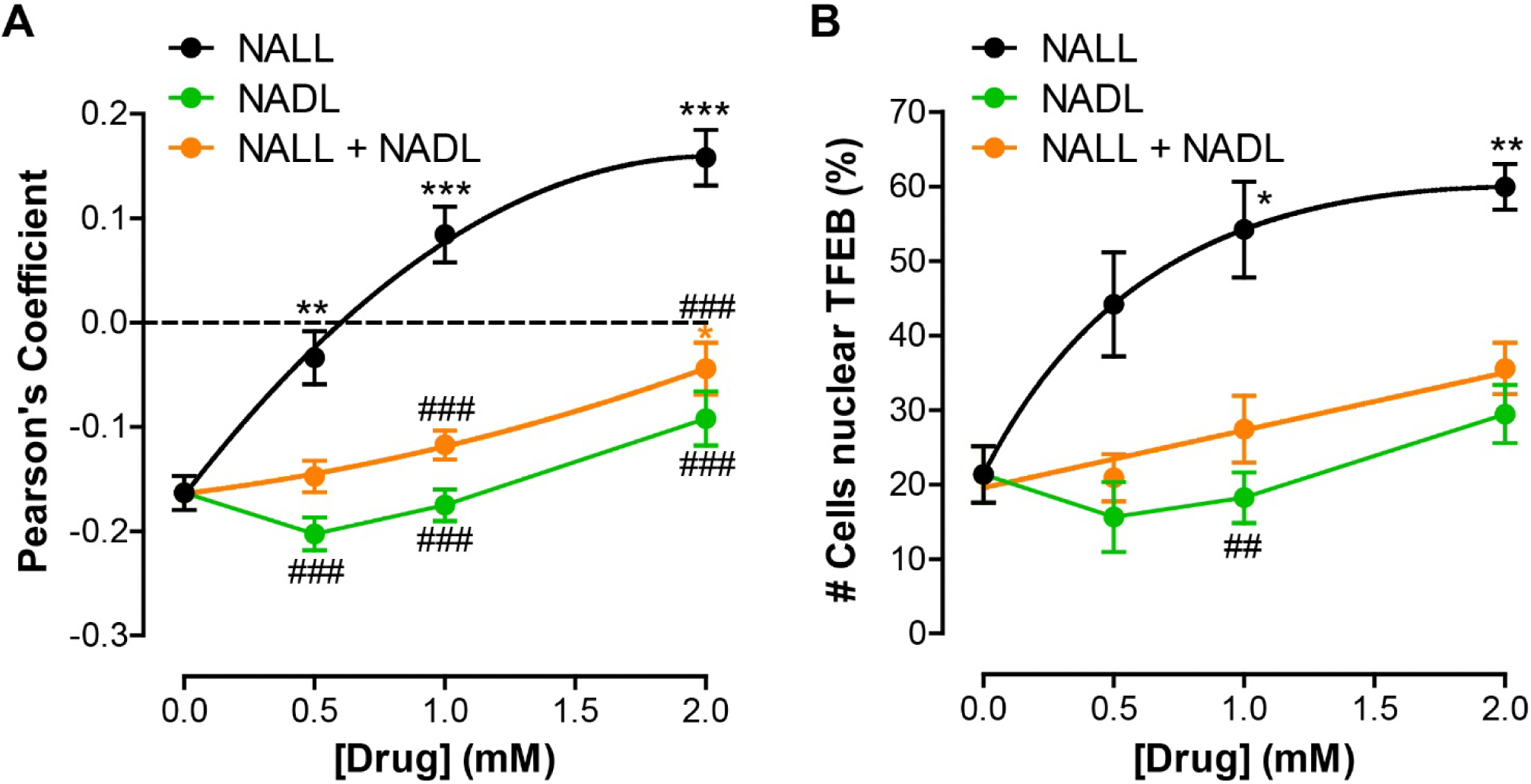
Levacetylleucine is the most efficacious enantiomer to evoke TFEB translocation. TFEB-EGFP translocation in HeLa cells incubated for 18 hours with different concentrations of each enantiomer: the L-form (levacetylleucine (NALL), black), D-form (N-acetyl D-leucine (NADL), green) or the racemic mixture of both (N-acetyl D-L-leucine (NALL + NADL), orange). Translocation is expressed either as the Pearson’s correlation coefficient with NucSpot 650 (A) or as the percentage of cells with nuclear TFEB (B). Data are expressed as the mean ± SEM of 153-265 cells, and significance determined by nonparametric ANOVA test, with significance denoted by * *p* < 0.05, ** *p* < 0.01, or *** *p* < 0.001 compared to vehicle control, i.e. 0 mM. Comparing NALL and NADL or racemic mixture, using a nonparametric ANOVA test: ##, *p* < 0.01; ###, *p* < 0.001.

Finally, we demonstrated that there is a downstream functional correlate of TFEB activation. As a master transcription factor, TFEB regulates lysosomal protein expression and lysosomal mass (Settembre et al., 2012). A major lysosomal marker protein is lysosome-associated membrane protein 1 (LAMP1), a transmembrane protein expressed in lysosomes and late endosomes. Here we show that incubation of cells with NALL significantly increased the expression of LAMP1 (**Fig. 4**). The LAMP1 gene has a CLEAR sequence in its promoter region, which affords regulation of its expression by TFEB.

**Figure 4.**
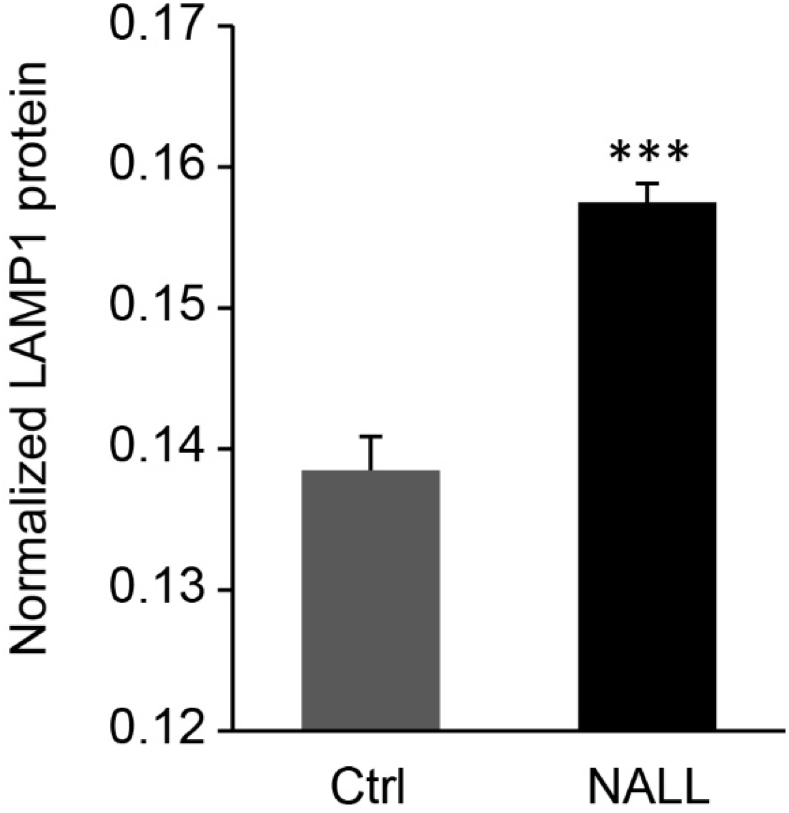
Levacetylleucine increases LAMP1 protein expression. HeLa cells were treated with 2 mM levacetylleucine (NALL) for 18 hours, then processed for In-Cell Western analysis: endogenous LAMP1 was labelled with an anti-LAMP1 antibody and stained with CellTag700 (for normalization to cell number). Data are expressed as the mean (LAMP1/CellTag700) ± SEM of 12 replicates, and significance determined by a paired t-test, with significance denoted by *** *p*< 0.001 compared to vehicle control (Ctrl), i.e. 0 mM NALL.

## Discussion

In this study, we further elucidated an aspect of NALL’s poly-pharmacological mechanism of action, demonstrating that the compound stimulates the translocation of TFEB to the nucleus and has a direct impact on lysosomal function. These findings are summarized in a schematic (**Fig. 5**) which illustrates the key actions of NALL and key signalling pathways it modulates, which are reflected in the changes in cellular responses seen. Importantly, this effect was not seen with non-acetylated L-leucine, nor the D enantiomer of N-acetyl leucine, demonstrating the potential of NALL to have this additional mechanism of action.

**Figure 5.**
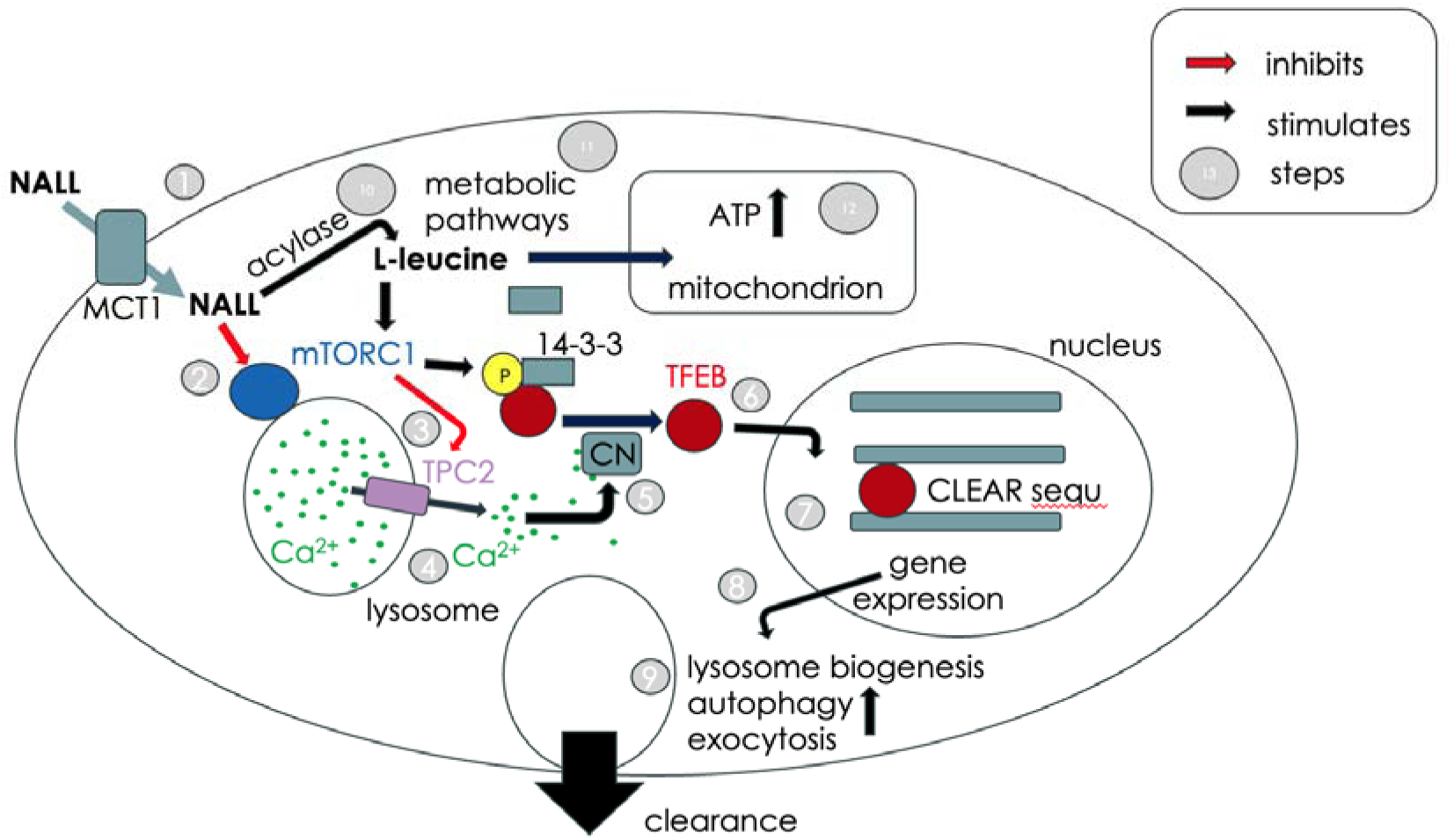
Proposed scheme for the mechanisms of action of levacetylleucine (NALL) to enhance lysosomal and mitochondrial function Step 1. Levacetylleucine (NALL) is transported into the cytosol of cells via the MCT transporter family. **Step 2** Levacetylleucine inhibits the activity of the mTORC complex and reduces TFEB phosphorylation. **Step 3** Inhibition of mTOR relives its inhibition on Ca^2+^ release channels expressed in lysosomes leading to Ca^2+^ release by TPC2 and/or TRPML1 (has this been shown as if not suggest proposed or hypothetical in title of Figure legend) (**Step 4**). **Step 5** Lysosomal Ca^2+^ release activates calcineurin (CN) to dephosphorylate TFEB and remove its ability to bind 14-3-3 proteins allowing it to translocate to the nucleus (**Step 6**). **Step 7** TFEB bind to CLEAR gene promoters and activates lysosomal and autophagic gene expression. **Step 8** This increases lysosomal biogenesis, autophagy and lysosomal exocytosis leading to cellular clearance of stored lysosomal material (**Step 9**). **Step 10** Levacetylleucine is metabolised to L-leucine that reactivates mTOR resulting in TFEB inactivation through phosphorylation. **Step 11** L-leucine enters metabolic pathways, where it increases mitochondrial ATP production (**Step 12**).

Previous studies have shown that NALL improves the bioenergetics of neurons by enhancing ATP production and activity of the citric acid cycle, as well as reducing lysosomal storage of cholesterol, sphingosine, and glycosphingolipids (GSLs) (Kaya et al., 2020). The efficacy of the drug is dependent on the acetyl moiety which renders the compound a substrate for high-capacity MCT transporters (Churchill et al., 2021), leading to high intracellular concentrations which are transformed into L-leucine by acylases, and have been shown to be expressed in HeLa cells. Such high concentrations of L-leucine would be expected to stimulate mTOR (Wolfson et al., 2016), and in consequence phosphorylate TFEB, reducing its transcriptional activity (as has been reported for another acetylated analogue of leucine, N-acetyl leucine amide (Hidayat et al., 2003)). It is hypothesized that this effect may be shared by NALL prior to its metabolism to L-leucine. However, in contrast, we find that NALL activates TFEB. A transient inhibition of mTOR may be sufficient to suppress mTOR-mediated TFEB phosphorylation to allow sufficient TFEB translocation to the nucleus to activate the CLEAR network of lysosomal and autophagy genes, as we have demonstrated for LAMP1 expression (**Fig. 4**) as an exemplar.

Importantly, the observed effects of NALL occur at concentrations which are in the human therapeutic range (Churchill et al., 2020). Specifically, our findings demonstrate that NALL (Aqneursa) at therapeutically relevant concentrations significantly enhances TFEB activation improving lysosomal biogenesis and function. TFEB activation has also recently been proposed as a mechanism of action for another newly approved drug for NPC, arimoclomol (Shammas et al., 2025), where an effect on TFEB activation was observed at much higher concentrations than for therapeutic doses (Cudkowicz et al., 2008).

Another finding from this study was that the effect of NALL was stereospecific in that N-acetyl D-leucine did not activate TFEB nuclear translocation (**Fig. 3**). The racemate, N-acetyl D, L-leucine was also without effect, indicating that the presence of equimolar N-acetyl D-leucine is antagonistic and inhibits the effect of the active L-enantiomer. Consistent with previous studies, NALL demonstrated considerable benefits over the racemate.

NALL’s ability to enhance TFEB nuclear translocation and lysosomal function provides further evidence of its potential utility as a treatment for a broad range of neurodegenerative and neurodevelopmental disorders characterized by lysosomal and dysfunction (Deus et al., 2020). For example, increases in TFEB have been suggested to enhance the degradation of Aβ plaques in mouse hippocampus by 40% (Xiao et al., 2015), which may be relevant for human disease since in preclinical models, a 25% Aβ reduction correlates with a meaningful improvement in cognitive deficits (Singh et al., 2024). It has also been shown that TFEB activation promotes the degradation of Tau proteins, leading to a reduction in Tau aggregates (Polito et al., 2014) and that a small molecule activator of TFEB ameliorates beta-amyloid precursor protein and Tau pathology in Alzheimer’s disease models (Song et al., 2016). Studies have confirmed that TFEB can reduce the deposition of toxic Tau aggregates, which represents a neuroprotective effect. TFEB enhances the lysosomal degradation of Tau, thereby reducing the formation of toxic aggregates (Cortes and La Spada, 2019) These mechanisms are crucial for the prevention of tauopathies, such as those observed in progressive supranuclear palsy and other neurodegenerative diseases such as Alzheimer’s disease.

TFEB also helps regulate the secretion of abnormal Tau, which promotes cellular clearance and can prevent further neuronal loss, and has been shown to have a protective effect against α-synuclein aggregates, which are typically associated with Parkinson’s disease. Recently, NALL has been shown to decrease pathological pS129-alpha-synuclein in dopaminergic neurons from patients harboring GBA1 or LRRK2 mutations, or in the LRRK2 (R1441C) knock-in mouse model of Parkinson’s disease (Song et al., 2025). This was paralleled by improved dopamine-dependent motor learning deficits as well as significant increases in 13 lysosomal proteins including HEXA, CTSA, HEXB, LAMP2, M6PR and PLA2G15 (Song et al., 2025). TFEB promotes autophagy and the lysosomal degradation of alpha-Synuclein oligomers, which can reduce the toxic effects of these aggregations (Decressac et al., 2013). In animal models, TFEB overexpression resulted in nearly complete prevention of dopaminergic neuronal loss, indicating the neuroprotective effect of TFEB. Moreover, TFEB overexpression in a mouse model increases dopamine release by 20%, reverses the atrophy of dopaminergic neurons, and enhances protein biosynthesis through mTORC1 phosphorylation (Torra et al., 2018).

The effects of acetyl-leucine on an alpha-synucleinopathy, Parkinson’s disease and its prodromal stage, REM sleep behaviour disorder (RBD) were recently demonstrated in a clinical case series. Acetyl-leucine not only eradicated symptoms of RBD, but remarkably, reversed loss of striatal dopamine-transporter binding and stabilized pathological metabolic brain pattern (Oertel et al., 2024), preventing the phenotypic conversion of patients from prodromal to manifest Parkinson’s Disease. This effect is consistent with the effect of TFEB overexpression/activation demonstrated in animal models. However, excessive TFEB activation is associated with tumorigenesis (Chen et al., 2024) and hyper-activation of autophagic pathways leading to cell death, and indeed small molecule inhibitors of TFEB have been developed and shown to inhibit autophagy (Lin et al., 2023). In contrast, NALL likely only transiently activates TFEB due to its rapid metabolism to L-leucine (Churchill et al., 2020), mitigating against the effects of excessive TFEB stimulation, but can also normalize excessive TFEB activation in disease cells. In a recent study, remarkably NALL corrects defective ER-lysosomal membrane contact in NPC1 patient fibroblast cells (Kiraly, 2025) which are thought to be due to a deficit in NPC1 protein-mediated tethering to the ER (Hoglinger et al., 2019). Whether modulation of the TFEB pathway by NALL leads to alternative tethering mechanisms is a possibility.

Our new finding that NALL has a direct impact on lysosomal function by activating the TFEB pathway (scheme: Fig. 5) provides further insights into the unique, poly-pharmacological mechanism of action of this compound. By targeting the mitochondrial-lysosomal axis (Deus et al., 2020), NALL may serve to ameliorate not only lysosomal storage disorders but also neurodegenerative and neurodevelopmental disorders more generally.

## Abbreviations

ATP: adenosine triphosphate
CLEAR: network coordinated lysosomal expression and regulation network
ER: endoplasmic reticulum
FDA: Food and Drug Administration (USA)
INN: International Nonproprietary Names
LAT: L-type amino-acid transporter
LSD: lysosomal storage disease
MCT: monocarboxylate transporters
mTORC1: mammalian target of rapamycin complex 1
NPC: Niemann-Pick disease type C
NALL: N-acetyl L-leucine
MCS: membrane contact site(s)
RBD: REM sleep behaviour disorder
TFEB: Transcription Factor EB
USAN: United States Adopted Name

---

**Fig. S1.**
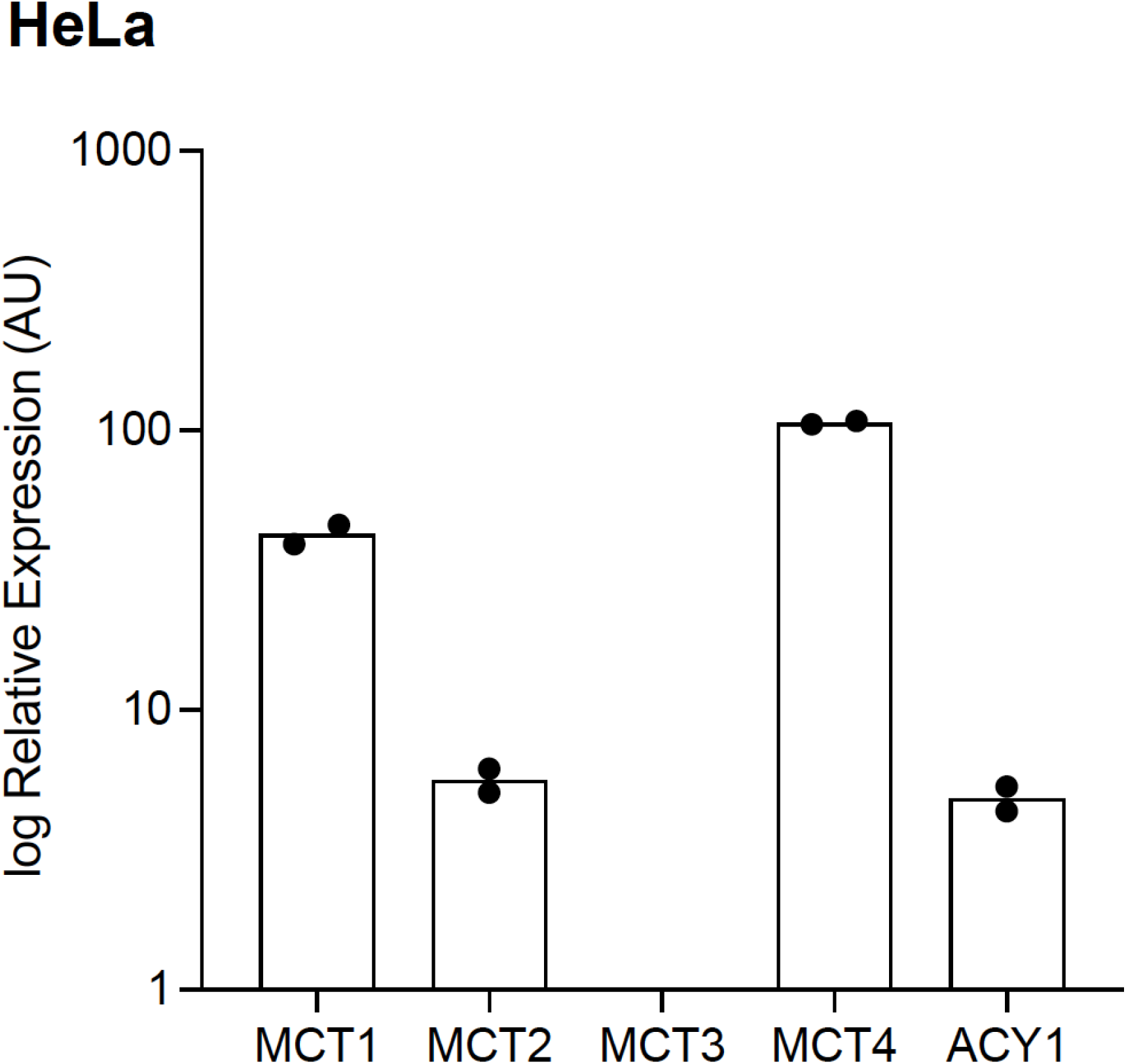
Expression patterns of monocarboxylate transporters (MCT1, MCT2, MCT3, MCT4) and the aminoacylase 1 enzyme (ACY1). Determined by RT-qPCR, expression was normalised to ß-actin, on a scale where ß-actin expression equals 10,000 arbitrary units. N=2.

